# Antagonistic coevolution between multiple quantitative traits: Matching dynamics can arise from difference interactions

**DOI:** 10.1101/509885

**Authors:** Masato Yamamichi, Kelsey Lyberger, Swati Patel

## Abstract

Coevolution is one of the major drivers of complex dynamics in population ecology. Historically, antagonistic coevolution in victim-exploiter systems has been a topic of special interest, and involves traits with various genetic architectures (e.g., the number of genes involved) and effects on interactions. For example, exploiters may need to have traits that “match” those of victims for successful exploitation (i.e., a matching interaction), or traits that exceed those of victims (i.e., a difference interaction). Different models exist which are appropriate for different types of traits, including Mendelian (discrete) and quantitative (continuous) traits. For models with multiple Mendelian traits, recent studies have shown that antagonistic coevolutionary patterns that appear as matching interactions can arise due to multiple difference interactions with costs of having large trait values. Here we generalize their findings to quantitative traits and show, analogously, that the multidimensional difference interactions with costs sometimes behave qualitatively the same as matching interactions. While previous studies in quantitative genetics have used the dichotomy between matching and difference frameworks to explore coevolutionary dynamics, we suggest that exploring multidimensional trait space is important to examine the generality of results obtained from one-dimensional traits.

## Introduction

Theories developed from specific models are initially only as general as their underlying models, until they are shown to hold in other modeling frameworks. Hence, it is critical to compare results from a variety of modeling frameworks to test their generality. Antagonistic coevolution in victim-exploiter systems (e.g., prey-predator or host-pathogen systems) has been intensively studied theoretically (Abrams, 2000; Nuismer, 2017). It is thought to be a major driver for the emergence and maintenance of biodiversity (Thompson, 2005) and feedbacks between interacting species can create complex ecological (e.g., Cortez and Weitz, 2014) and evolutionary (e.g., Dercole et al., 2010) dynamics. Historically, various coevolution models with differing genetic architectures (e.g., the number of genes involved) and interspecific interactions (e.g., a “matching” vs. a “difference” interaction: see below) have been proposed and analyzed (Nuismer, 2017). To account for genetic architecture, one approach is to consider coevolution with a few major genes that interact and produce a discrete phenotype (Mendelian trait-based models, or hereafter Mendelian models) (e.g., Mode, 1958; Seger, 1988). On the other hand, quantitative trait-based models (hereafter quantitative models) assume the genetic architecture consists of many genes of small effect and produce continuous phenotypes (e.g., Gavrilets, 1997; Saloniemi, 1993).

Here we highlight the connection between Mendelian and quantitative models of victims and exploiters, and then clarify the relationships between existing modeling frameworks for quantitative traits. We begin by reviewing the different modeling frameworks for Mendelian traits. We then draw analogies between Mendelian and quantitative models (e.g., Boots et al., 2014; Cortez and Weitz, 2014). The analogies suggest that the matching framework, in which exploitation is more successful when traits “match” (i.e., a bidirectional axis of vulnerability *sensu* Abrams 2000), can arise from the difference framework, in which exploitation is more successful when traits differ (i.e., a unidirectional axis of vulnerability *sensu* Abrams 2000), in multidimensional quantitative models with costs of having large trait values. Finally, as a proof of concept, we give an example model of theoretical quantitative genetics where a victim-exploiter coevolutionary dynamics arising from two pairs of difference traits can appear as a matching interaction in a single pair of traits.

## Background

Historically, in Mendelian models, two major interactions were often assumed and contrasted: the matching allele (MA) framework, in which exploitation is successful when traits match (e.g., Grosberg and Hart, 2000) (Figure 1a), and the gene-for-gene (GFG) framework, in which a generalist exploiter can exploit a range of victim genotypes, but pays a cost for generalism (Flor, 1956) (Figure 1c). Researchers have investigated the relative importance and relationships of the two interactions (e.g., Agrawal and Lively, 2002; Frank, 1996; Parker, 1994). Recently, Ashby and Boots (2017) found that, in multidimensional Mendelian (multilocus) models, the GFG framework can behave effectively as the MA framework (Figure 2a).

**FIGURE 1.**
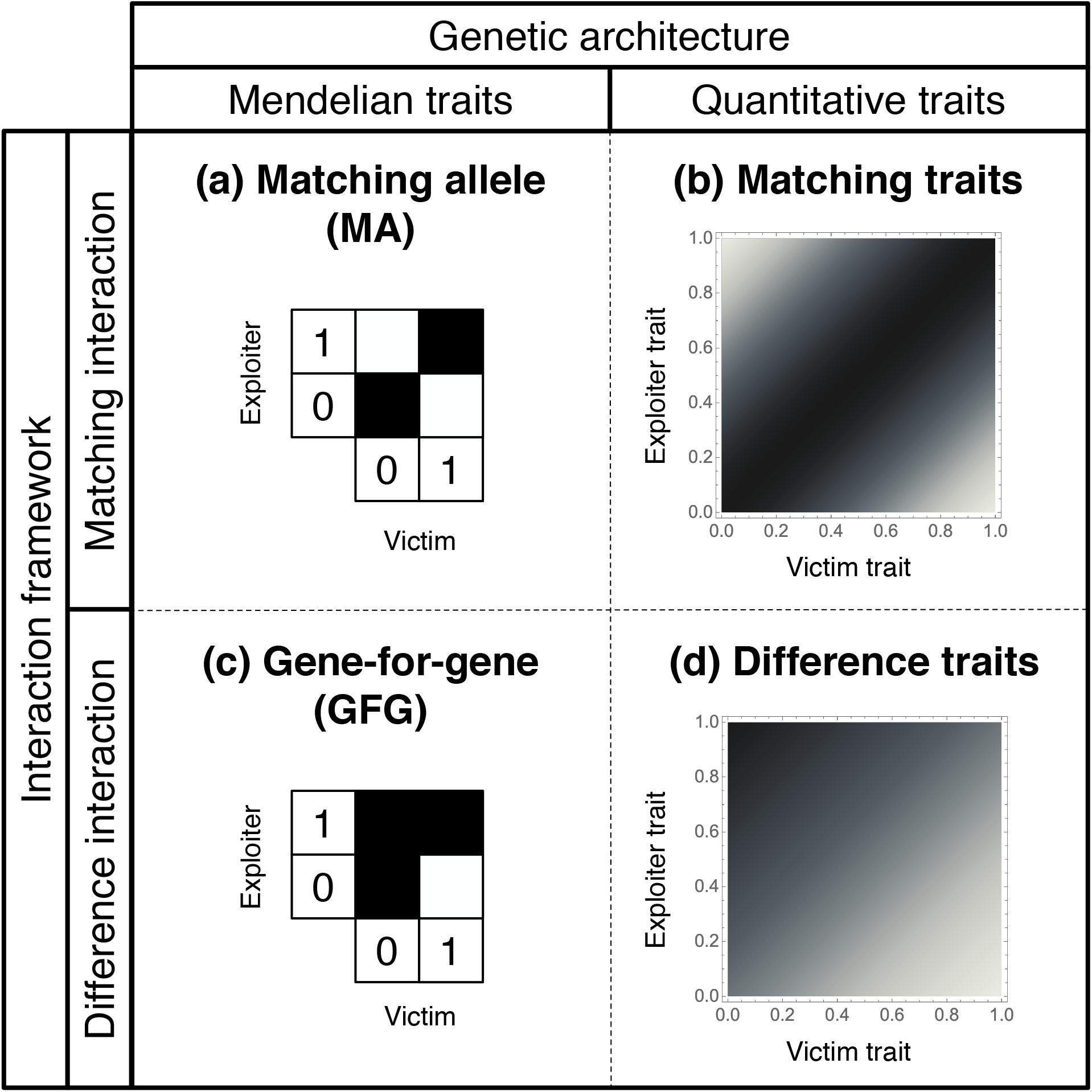
Interaction frameworks and genetic architectures in coevolution models. (a) Matching allele (MA) is a matching interaction in Mendelian traits. Victims and exploiters are haploid and their genotypes are represented by 1 and 0 at a particular locus. Dark shading indicates greater exploitation success. (b) The matching interaction is a MA interaction in quantitative traits. Exploitation success is maximized when the trait difference is zero. (c) Gene-for-gene (GFG) is a difference interaction in Mendelian traits. Here 1 and 0 indicate the presence and absence, respectively, of defense/exploitation alleles. (d) The difference interaction is a GFG interaction in quantitative traits. Exploitation success is increased by larger exploiter trait values and by smaller victim trait values.

**FIGURE 2.**
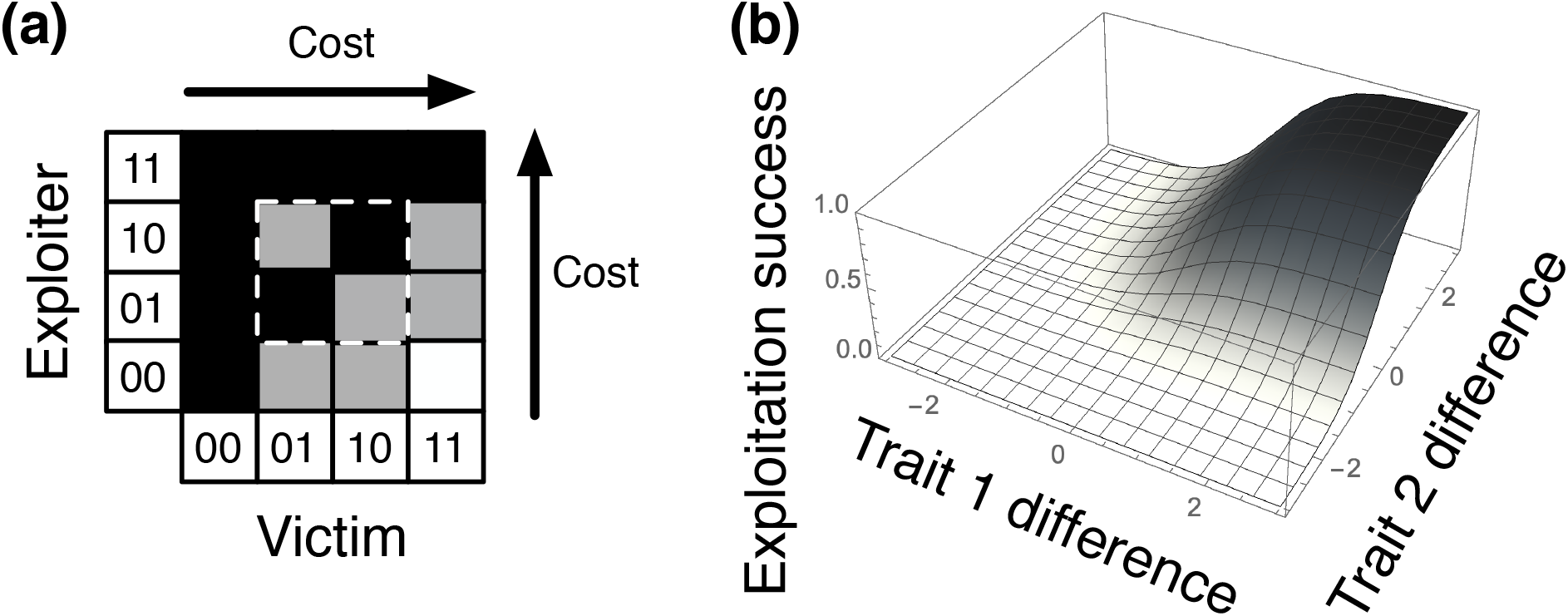
(a) An example of exploitation matrix in a Mendelian GFG model with two loci (Ashby and Boots, 2017). The region surrounded by a dashed white line shows that the MA interaction arises from the GFG framework. Victims and exploiters are haploid and their genotypes are represented by 1 and 0 at a locus, and they indicate the presence and absence, respectively, of defense/exploitation alleles. Darker shading indicates greater exploitation success and arrows indicate greater costs. (b) Exploitation success as a function of both trait differences between exploiters and victims (*y*_*i*_ − *x*_*i*_) in a quantitative difference model with two traits (Equation 1).

To illustrate, consider a simple GFG model with haploid victims and exploiters with two loci with two alleles each, labelled 1 and 0. The alleles 1 and 0 indicate the presence and absence, respectively, of an exploitation or defense trait (Figure 2a). If both of the allelic values of the exploiter are greater than or equal to those of the victim, maximum exploitation will occur (black squares in Figure 2a). If the victim has exactly one allelic value greater than that of the exploiter, so that the other allelic value of the victim is less than or equal to the corresponding value of the exploiter, then exploitation will be partial (gray squares in Figure 2a). Finally, in the case where both allelic values of the victim are strictly greater than that of the exploiter (which only occurs when the victim has genotype 11 and the exploiter has 00), then the victim is defended against exploitation (a white square in Figure 2a).

When coevolutionary dynamics occurs between four genotypes with the alleles 1 and 0, the type of interaction (indicated by a dashed white line in Figure 2a) effectively reduces to a MA interaction: the exploiter genotype 01 (10) is more successful at exploiting its matching counter victim genotype 01 (10) as in Figure 1a. Indeed, by investigating a multilocus GFG model, Ashby and Boots (2017) found that, in the case where an intermediate number of alleles is optimal, the victim/exploiter genotypes alternate between various subsets of defense/exploitation alleles, but the total number of alleles stays constant. Hence, the multidimensional GFG framework effectively behaves as the MA framework (Figure 2a).

As with Mendelian models, quantitative models have assumed an apparent dichotomy between two frameworks (Abrams, 2000; Abrams and Matsuda, 1997; Gilman et al., 2012; McPeek, 2017): the matching trait framework, in which exploitation is increasingly more successful with the extent that trait values of exploiters match those of victims (Figure 1b), and the difference trait framework, in which exploitation is increasingly more successful with the extent that trait values of exploiters exceed those of victims (Figure 1d). They are also known as the bidirectional and unidirectional trait cases, respectively (Abrams, 2000). The matching framework is analogous to the MA model and is used, for example, when an exploiter can best exploit a victim with body size or color within a certain range (Brown and Vincent, 1992; Calcagno et al., 2010; Dercole et al., 2010; Dieckmann et al., 1995; Fleischer et al., 2018; Gavrilets, 1997; Khibnik and Kondrashov, 1997; Kopp and Gavrilets, 2006; Marrow et al., 1992; Mougi, 2012; Nuismer et al., 2005; Yamamichi and Ellner, 2016). This has been observed in the coevolution between the African tawny-flanked prinia and its brood parasite, the cuckoo finch: the parasite varies its egg color to match that of the victim (Spottiswoode and Stevens, 2012). On the other hand, the difference framework is analogous to the GFG model and is appropriate when exploitation success is determined by speed vs. speed, strength vs. armor, or toxicity vs. resistance (Cortez and Weitz, 2014; Frank, 1994; Mougi and Iwasa, 2010; Mougi and Iwasa, 2011; Northfield and Ives, 2013; Nuismer et al., 2007; Saloniemi, 1993; Sasaki and Godfray, 1999; Tien and Ellner, 2012; van Velzen and Gaedke, 2017; Yamamichi and Miner, 2015). For example, garter snakes evolve stronger resistance to rough-skinned newt toxicity in areas where newts are more toxic (Brodie III and Brodie Jr, 1990).

In the matching framework, exploitation success is typically a unimodal function of the trait difference, and is often modeled as a Gaussian: exp[−(*x* − *y*)^2^], where *x* and *y* are traits of victims and exploiters, respectively (Figure 1b) (e.g., Gavrilets, 1997; Khibnik and Kondrashov, 1997; Mougi, 2012; Nuismer et al., 2005). On the other hand, in the difference framework, exploitation success is often modeled as a sigmoidal function of the trait difference: 1/[1 + exp(*x* − *y*)] (Figure 1d) (e.g., Mougi and Iwasa, 2010; Nuismer et al., 2007; van Velzen and Gaedke, 2017). In this framework, it is always advantageous for both species to increase the trait values for better exploitation/defense (McPeek, 2017). Empirical evidence suggests that there is an energetic trade-off to exploitation and defense and hence, most models of difference traits assume there is a cost associated with each trait, which prevents runaway evolution. For example, garter snakes resistant to newt toxin have costs associated with slower sprinting speeds (Brodie III and Brodie Jr, 1999), milkweed bugs’ production of enzymes to break down milkweed toxins has metabolic costs (Dalla and Dobler, 2016), and numerous plants produce defense chemicals with a cost to plant growth (reviewed in Herms and Mattson, 1992).

Here, by drawing analogies between the Mendelian trait and quantitative trait frameworks (Figure 1), we further extend the generalization of Ashby and Boots (2017) and demonstrate that the difference framework can result in similar dynamics to the matching framework in multidimensional quantitative models. The matching model is a continuous trait analog to the MA model and the difference model is a continuous trait analog to the GFG model (Figure 1). In a multilocus GFG model, under certain conditions the victim/exploiter genotypes alternate between various subsets of defense/exploitation alleles, but the total number of alleles stays constant, which is effectively what occurs under the MA framework (Ashby & Boots 2017). This is analogous to a bivariate difference trait model, in which the victim/exploiter genotypes alternate between having more investment in one defense/exploitation trait vs. another, which is effectively what occurs under the matching framework. Instead of switching between the extremes of a single matching trait, the species are switching between two difference-trait-based strategies. We consider a victim-exploiter model in which each has two difference traits that influence exploitation success. An empirical example that motivates our study is a stepwise infection process (Hall et al., 2017), where host species prevent exploitation of parasites by several difference traits. In the case where intermediate trait values are optimal due to the costs of defense/offense, the victim/exploiter traits may switch between combinations where a trait is large while the other trait is small, but the total costs stay constant. The resultant dynamics may appear as the matching interaction, and this is the quantitative-model analog to the findings in Ashby and Boots (2017) in Mendelian models.

## AN EXAMPLE

We develop a two-dimensional quantitative model with difference interactions (Figure 2b) to demonstrate that matching dynamics can emerge from a multidimensional difference framework (see Appendix S1 and Supplementary Mathematica file, Supporting Information, for details). We consider a quantitative genetic model (Iwasa et al., 1991; Lande, 1976) in continuous time (i.e., Malthusian fitness) (Abrams et al., 1993). We assume that victims and exploiters each have two quantitative traits. We let *x*_*i*_ be the value of victim trait *i* (= 1, 2) and assume greater values allow for defense against trait *i* of the exploiter. We let *y*_*i*_ be the value of exploiter trait *i* and assume greater values allow for more exploitation. Each trait is associated with a cost that decreases individual fitness with greater trait values. The difference between the exploiter trait and the corresponding victim trait, *y*_*i*_ − *x*_*i*_, affects exploitation success (Figure 2b), which is expressed by a sigmoid function (Figure 1d). We assume that fitness of victims and exploiters are, respectively,

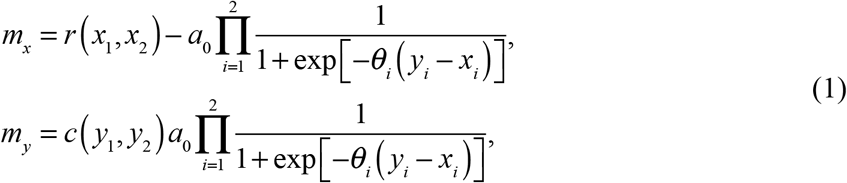

where *a*_0_ is the effects of successful exploitation on fitness (e.g., maximum attacking rate), and *θ*_*i*_(*i* = 1, 2) determine the sensitivity of exploitation success to the trait difference. In our model, the product 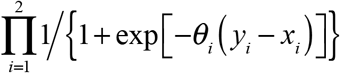 means that victims need to overcome their exploiters at only one trait for escaping from exploitation, whereas exploiters must overcome the two defense traits for successful exploitation (Gilman et al. 2012). This sequential interaction is also found in many host-parasite systems (Hall et al. 2017).

Here *r* and *c* represent costs of defense and exploitation for victims and exploiters, respectively (e.g., prey growth and predator conversion efficiency are decreasing functions of defense and offense, respectively: see Appendix S1, Supporting Information), and they are linearly decreasing functions of the sum of the two traits values (i.e., lower values of *r* and *c* represent higher costs). The costs can prevent escalation toward positive infinite values in the two traits. However, the two-dimensional difference model with costs behaves like a one-dimensional matching model, in which runaway evolution occurs toward a positive infinite value in one trait and a negative infinite value in the other trait (Appendices S2, S3, Figures S1, S2, Supporting Information). To prevent this, we add stabilizing selection to the victim cost function as in Gavrilets (1997). This can occur in quantitative models but not in Mendelian models because it is customary to assume unbounded trait values in quantitative models, which permits the existence of runaway escalation, whereas Mendelian traits are typically modeled as taking values in finite sets. We employ the following functions:

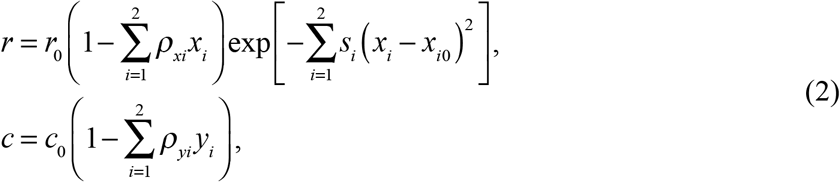

where *r*_0_ and *c*_0_ are basic parameter values (e.g., for prey growth rate and predator conversion efficiency) and *ρ*_*ij*_ (*i* = *x*, *y*, *j* = 1, 2) determine the slope of the functions. Stabilizing selection drives victim evolution toward (*x*_1_, *x*_2_) = (*x*_10_, *x*_20_), where *s*_*i*_ (*i* = 1, 2) determines the strength of stabilizing selection (Figure S3, Supporting Information). Note that adding stabilizing selection to the exploiter function instead of the victim function results in victim’s escape from exploitation. We can add stabilizing selection to the exploiter function in addition to the victim function, but here we keep the model as simple as possible.

Then coevolutionary dynamics are described by four ordinary differential equations,

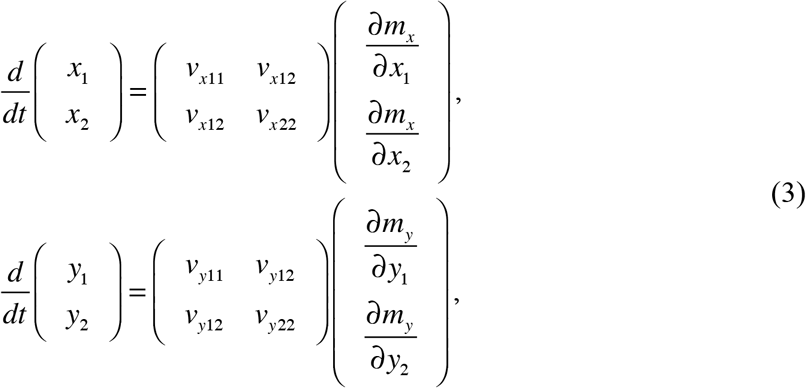

where *v*_*ijj*_ (*i* = *x*, *y*, *j* = 1, 2) represents additive genetic variance, *v*_*i*12_ (*i* = *x*, *y*) represents covariance, and partial derivatives represent fitness gradients (Abrams et al., 1993). For simplicity, we assume that the traits evolve independently (genetic covariance, *v*_*i*12_, is zero). For example, this is likely in hosts that defend against parasites by a cellular immune response and a behavioral response, but further work is needed to explore the effects of trait correlations. Also, our trait symmetry assumptions (i.e., the additive genetic variances are the same, *v*_*i*11_ = *v*_*i*22_, and the two traits affect exploitation success and fitness costs in the same way, *θ*_1_ = *θ*_2_, *ρ*_*x*1_ = *ρ*_*x*2_, *ρ*_*y*1_ = *ρ*_*y*2_, and *s*_1_ = *s*_2_) should be carefully examined in future studies as this assumption may not hold for most traits (Figure S4, Supporting Information), and it may result in, for example, time-scale separation in evolutionary dynamics of the two traits.

By numerically solving the model, we find that the two-dimensional difference model can behave as a one-dimensional matching model. First, when there is no stabilizing selection in the victim cost (*s*_*i*_ = 0) and the exploiter genetic variance is small enough (*v*_*xii*_ >> *v*_*yii*_, *i* = 1, 2), coevolutionary dynamics shows runaway escalation toward extreme trait values, and the coevolutionary outcome depends on initial conditions (Figure S2, Supporting Information). Second, when there is stabilizing selection (*s*_*i*_ > 0) and the exploiter genetic variance is large enough (e.g., *v*_*xii*_ < *v*_*yii*_), the system is attracted to a stable equilibrium (the blue and red points in Figures 3a, 3b), and there is disruptive selection for victim traits (i.e., fitness minimum: vectors representing fitness gradients in Figure 3a are distracted from the equilibrium) and stabilizing selection for exploiter traits (i.e., fitness maximum: vectors in Figure 3b are attracted to the equilibrium), which is consistent with the matching interaction (Appendix S2, Supporting Information) (Abrams and Matsuda, 1997; Gavrilets, 1997). Third, when there is stabilizing selection (*s*_*i*_ > 0) and the exploiter genetic variance is small enough (*v*_*xii*_ >> *v*_*yii*_), the equilibrium is not stable (Appendix S3, Figure S3, Supporting Information) and the trait dynamics fluctuate (Figure 3c) but are constrained to regions in which the total investments are nearly constant (Figure 3d) and the two traits for each species show antiphase cycles (Figures 3c, 3e, 3f). These simulation results demonstrate that the matching dynamics can arise from the difference interaction, generalizing Ashby and Boots (2017)’s idea from Mendelian models to quantitative models.

**FIGURE 3.**
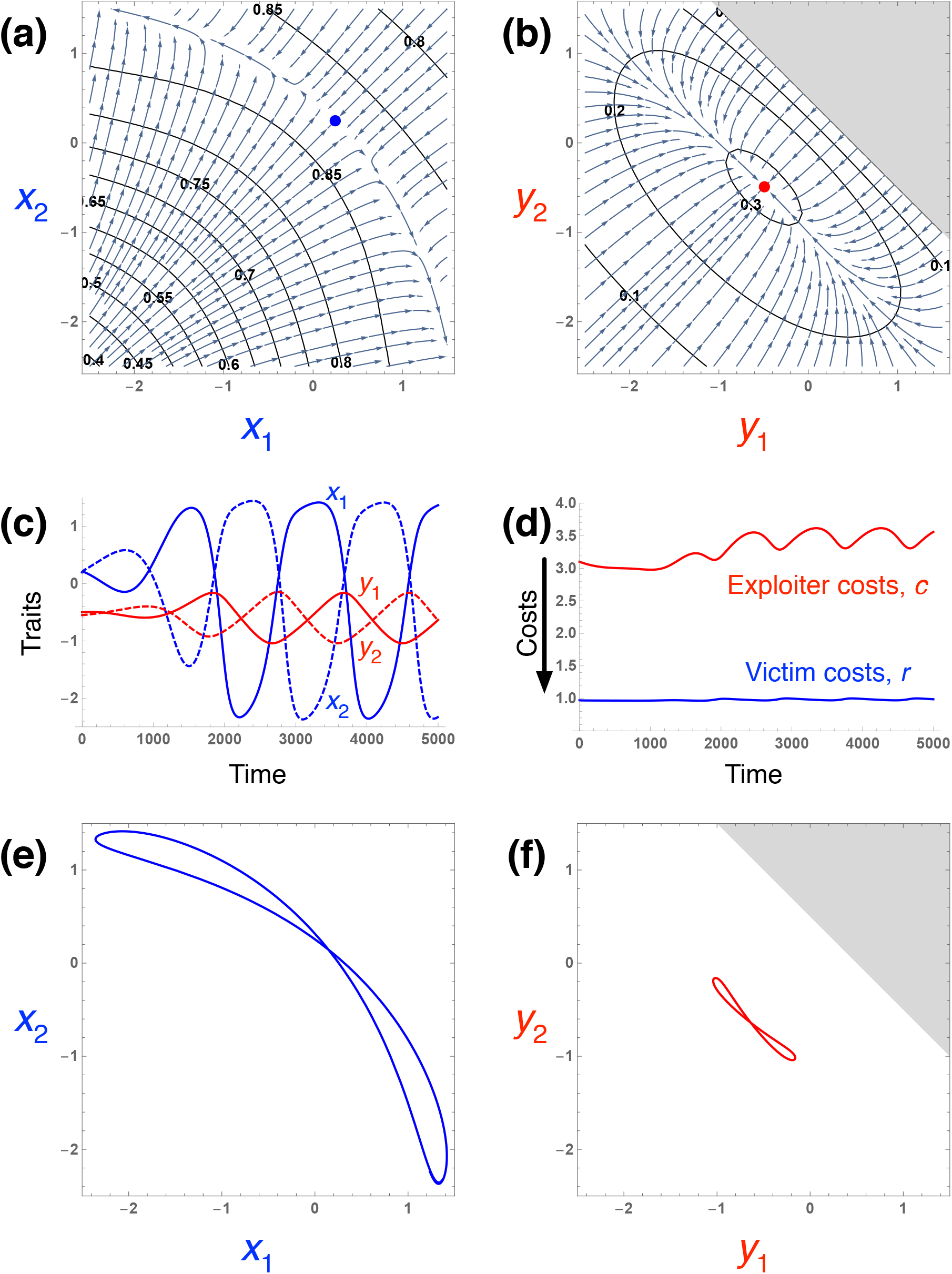
Coevolutionary dynamics of two quantitative traits with difference interactions. (a, b) Fitness landscapes of victims (a) and exploiters (b) when the exploiter genetic variance is large enough and the system converges to a stable equilibrium. (a) A contour plot of fitness (*m*_*x*_) and a vector field of fitness gradients for the victim traits when the exploiter traits are at the equilibrium trait value (the red point in b). (b) A contour plot of fitness (*m*_*y*_) and a vector field of fitness gradients for the exploiter traits when the victim traits are at the equilibrium trait value (the blue point in a). Parameter values are *ρ*_*xi*_ = 0.07, *ρ*_*yi*_ = 2, *x*_10_ = *x*_20_ = 0.2, *a*_0_ = *r*_0_ = *c*_0_ = *θ*_*i*_ = *v*_*jii*_ = 1, *s*_*i*_ = 0.01, and *v*_*j*12_ = 0 (*i* = 1, 2, *j* = *x*, *y*). (c-f) Coevolutionary cycles when the victim genetic variance is much larger than exploiter genetic variance (*v*_*xii*_ >> *v*_*yii*_). (c) Numerical solutions of the four traits. (d) Numerical solutions of the functions (Equation 2) representing costs of defense/exploitation (note that lower values indicate higher costs). (e, f) Coevolutionary cycles of victims (e) and exploiters (f). Parameter values are the same as (a) and (b) except for *v*_*xii*_ = 1 and *v*_*yii*_ = 0.01 (*i* = 1, 2). Note that *c* is negative in gray regions in (b) and (f).

## Discussion

Victim-exploiter interactions are among the most fundamental type of ecological interactions. In addition to their importance in ecological communities, these interactions are widely recognized for playing an important role in ecological dynamics (e.g., extinction and predator-prey cycles: Cortez and Weitz (2014); Northfield and Ives (2013)) and evolutionary dynamics (e.g., sex, recombination, and epistasis: Otto and Lenormand (2002)), because each species needs to continue to adapt to new selection pressures by the other species. Due to its importance, many modeling frameworks have been proposed to investigate the dynamics of coevolution between a victim and its exploiter, capturing various aspects of the traits involved (Nuismer, 2017).

In this work, we highlight the relationship between two modeling frameworks of quantitative traits in victim-exploiter coevolution. In particular, a multidimensional difference trait framework, in which each trait confers an advantage in the victim-exploiter interaction but is associated with some cost (Figures 1d, 2b) can generate dynamics like a matching trait framework, in which traits between a victim and exploiter must “match” in some sense for successful interactions (Figure 1b). Our investigation extends our knowledge of the coevolutionary process by demonstrating how effectively matching dynamics can be generated from difference traits with costs in a special case of multivariate coevolution as a proof of concept (rather than a thorough investigation of the dynamics: Figure 3).

For some traits, the distinction between matching traits and difference traits is clear. For both victims and exploiters, being stronger and faster are always more advantageous (but may have an associated cost or energetic limitation) and hence, are difference traits. However, in other cases, it is not so obvious. For example, body size of both victims and exploiters can be a difference trait, in which bigger is better for exploitation/defense (e.g., gape-limited predation), or a matching trait, in which exploiters must be within a certain size range depending on victims’ size for successful exploitation. Besides classifying difference vs. matching traits, a second empirical difficulty arises when not all traits are simultaneously measured. Our simulations suggest that, even when the body size is a difference trait, it may behave as a matching trait because of another potentially unmeasured difference trait such as toxicity/resistance. This correlated coevolution between the two traits can occur even without genetic covariance, as long as the body size and toxicity/resistance affect exploitation success and the costs of being big affect fitness together with the costs of the toxicity/resistance.

Previous work has suggested that matching traits are more likely to lead to coevolutionary cycling of traits than difference interactions (Abrams, 2000; McPeek, 2017). In comparing quantitative models with a one-dimensional trait, for example, McPeek (2017) found cycles are observed in a smaller area of parameter space with difference traits compared to matching traits. Furthermore, the underlying mechanism of cycles differs in the two frameworks. In the matching framework, cycles consist of the exploiter tracking victim traits and the victim escaping from exploiter traits (i.e., fluctuating selection between equally specific defense/exploitation traits). In the difference framework, cycles consist of the victim and exploiter investing more energy in their traits for better defense/exploitation, and then abandoning their defense/exploitation because at some point they are too costly to maintain (i.e., fluctuating selection between strong and weak defense/exploitation traits). Even when empirical researchers are focusing on a single trait with the difference interaction, the system may have another difference trait that can affect exploitation success. In such a case, coevolutionary dynamics can be driven by the matching-like interaction, which may cause cyclic dynamics more easily (McPeek, 2017).

In spite of recent interest in coevolution in multidimensional trait space (e.g., Débarre et al., 2014; Doebeli and Ispolatov, 2017; Gilman et al., 2012), the relation of the matching and difference frameworks has not been well recognized. For example, in their study on evolutionary escape, Gilman et al. (2012) contrasted coevolutionary dynamics with matching and difference interactions in multidimensional trait space but did not find a significant difference between them (see Figure 1 of Gilman et al. (2012)). Our generalization offers a potential explanation for this previous work, as coevolutionary dynamics with matching interactions can arise from difference interactions in multidimensional quantitative models.

This work was inspired by the recent work of Ashby and Boots (2017), who showed an analogous relationship for two Mendelian trait models of coevolution. Here, we draw out the parallels between two Mendelian (single-locus) trait frameworks and two quantitative trait frameworks. Namely, we highlight that the matching allele (MA) framework is a Mendelian analog of the matching framework, and the gene-for-gene (GFG) framework is a Mendelian analog of the difference framework (Figure 1). While these frameworks are valuable, many traits important to victim-exploiter interactions may actually lie somewhere in the middle of these two extremes; they may be determined by many (but not infinite number of) loci. It is still not well understood how coevolutionary dynamics differ between when traits are governed by a single locus, multi-locus (with linkage disequilibrium and epistasis), and effectively infinite loci (as assumed in classical quantitative genetics). While there have been a handful of papers to investigate this question in specific scenarios, further investigation is need for a general understanding.

## Acknowledgements

We thank two anonymous reviewers, P. A. Abrams, M. H. Cortez, B. Ashby, and M. Boots for helpful comments. M.Y. was supported by the Japan Society for the Promotion of Science (JSPS) Grant-in-Aid for Scientific Research (KAKENHI) 15H02642, 16K18618, 16H04846, 18H02509, and 19K16223, and by John Mung Program and Hakubi Center for Advanced Research of Kyoto University. K.L. was supported by National Science Foundation (NSF) Grant DMS-1313418 to S. J. Schreiber at the University of California, Davis. S.P. was supported by the Austrian Science Fund (FWF) Grant P25188-N25 to R. Bürger at the University of Vienna.

## REFERENCES

Abrams, P. A. (2000) The evolution of predator-prey interactions: Theory and evidence. Annual Review of Ecology and Systematics 31: 79–105. DOI 10.1146/annurev.ecolsys.31.1.79.

Abrams, P. A. & Matsuda, H. (1997) Fitness minimization and dynamic instability as a consequence of predator-prey coevolution. Evolutionary Ecology 11: 1–20. DOI 10.1023/a:1018445517101.

Abrams, P. A., Matsuda, H. & Harada, Y. (1993) Evolutionarily unstable fitness maxima and stable fitness minima of continuous traits. Evolutionary Ecology 7: 465–487. DOI 10.1007/BF01237642.

Agrawal, A. & Lively, C. M. (2002) Infection genetics: gene-for-gene versus matching-alleles models and all points in between. Evolutionary Ecology Research 4: 79–90.

Ashby, B. & Boots, M. (2017) Multi-mode fluctuating selection in host–parasite coevolution. Ecology Letters 20: 357–365. DOI 10.1111/ele.12734.

Boots, M., White, A., Best, A. & Bowers, R. (2014) How specificity and epidemiology drive the coevolution of static trait diversity in hosts and parasites. Evolution 68: 1594–1606. DOI 10.1111/evo.12393.

Brodie III, E. D. & Brodie Jr, E. D. (1990) Tetrodotoxin resistance in garter snakes: an evolutionary response of predators to dangerous prey. Evolution 44: 651–659. DOI 10.1111/j.1558-5646.1990.tb05945.x.

Brodie III, E. D. & Brodie Jr, E. D. (1999) Costs of exploiting poisonous prey: evolutionary trade-offs in a predator-prey arms race. Evolution 53: 626–631. DOI 10.1111/j.1558-5646.1999.tb03798.x.

Brown, J. S. & Vincent, T. L. (1992) Organization of predator-prey communities as an evolutionary game. Evolution 46: 1269–1283. DOI 10.2307/2409936.

Calcagno, V., Dubosclard, M. & de Mazancourt, C. (2010) Rapid exploiter-victim coevolution: The race is not always to the swift. The American Naturalist 176: 198–211. DOI 10.1086/653665.

Cortez, M. H. & Weitz, J. S. (2014) Coevolution can reverse predator–prey cycles. Proceedings of the National Academy of Sciences of the United States of America 111: 7486–7491. DOI 10.1073/pnas.1317693111.

Dalla, S. & Dobler, S. (2016) Gene duplications circumvent trade-offs in enzyme function: Insect adaptation to toxic host plants. Evolution 70: 2767–2777. DOI 10.1111/evo.13077.

Débarre, F., Nuismer, S. L. & Doebeli, M. (2014) Multidimensional (co)evolutionary stability. The American Naturalist 184: 158–171. DOI 10.1086/677137.

Dercole, F., Ferriere, R. & Rinaldi, S. (2010) Chaotic Red Queen coevolution in three-species food chains. Proceedings of the Royal Society B: Biological Sciences 277: 2321–2330. DOI 10.1098/rspb.2010.0209.

Dieckmann, U., Marrow, P. & Law, R. (1995) Evolutionary cycling in predator-prey interactions: population dynamics and the Red Queen. Journal of Theoretical Biology 176: 91–102. DOI 10.1006/jtbi.1995.0179.

Doebeli, M. & Ispolatov, I. (2017) Diversity and coevolutionary dynamics in high-dimensional phenotype spaces. The American Naturalist 189: 105–120. DOI 10.1086/689891.

Fleischer, S. R., terHorst, C. P. & Li, J. (2018) Pick your trade-offs wisely: Predator-prey eco-evo dynamics are qualitatively different under different trade-offs. Journal of Theoretical Biology 456: 201–212. DOI 10.1016/j.jtbi.2018.08.013.

Flor, H. H. (1956) The complementary genic systems in flax and flax rust. Advances in genetics 8: 29–54. DOI 10.1016/S0065-2660(08)60498-8.

Frank, S. A. (1994) Coevolutionary genetics of hosts and parasites with quantitative inheritance. Evolutionary Ecology 8: 74–94. DOI 10.1007/BF01237668.

Frank, S. A. (1996) Problems inferring the specificity of plant-pathogen genetics. Evolutionary Ecology 10: 323–325. DOI 10.1007/BF01237689.

Gavrilets, S. (1997) Coevolutionary chase in exploiter-victim systems with polygenic characters. Journal of Theoretical Biology 186: 527–534. DOI 10.1006/jtbi.1997.0426.

Gilman, R. T., Nuismer, S. L. & Jhwueng, D.-C. (2012) Coevolution in multidimensional trait space favours escape from parasites and pathogens. Nature 483: 328–330. DOI 10.1038/nature10853.

Grosberg, R. K. & Hart, M. W. (2000) Mate selection and the evolution of highly polymorphic self/nonself recognition genes. Science 289: 2111–2114. DOI 10.1126/science.289.5487.2111.

Hall, M. D., Bento, G. & Ebert, D. (2017) The evolutionary consequences of stepwise infection processes. Trends in Ecology & Evolution 32: 612–623. DOI 10.1016/j.tree.2017.05.009.

Herms, D. A. & Mattson, W. J. (1992) The dilemma of plants: to grow or defend. The Quarterly Review of Biology 67: 283–335. DOI 10.1086/417659.

Iwasa, Y., Pomiankowski, A. & Nee, S. (1991) The evolution of costly mate preferences II. The ‘handicap’ principle. Evolution 45: 1431–1442. DOI 10.2307/2409890.

Khibnik, A. I. & Kondrashov, A. S. (1997) Three mechanisms of Red Queen dynamics. Proceedings of the Royal Society B: Biological Sciences 264: 1049–1056. DOI 10.1098/rspb.1997.0145.

Kopp, M. & Gavrilets, S. (2006) Multilocus genetics and the coevolution of quantitative traits. Evolution 60: 1321–1336. DOI 10.1111/j.0014-3820.2006.tb01212.x.

Lande, R. (1976) Natural selection and random genetic drift in phenotypic evolution. Evolution 30: 314–334. DOI 10.2307/2407703.

Marrow, P., Law, R. & Cannings, C. (1992) The coevolution of predator-prey interactions: ESSs and Red Queen dynamics. Proceedings of the Royal Society B: Biological Sciences 250: 133–141. DOI 10.1098/rspb.1992.0141.

McPeek, M. A. (2017) Evolutionary Community Ecology. Princeton University Press, Princeton, NJ.

Mode, C. J. (1958) A mathematical model for the co-evolution of obligate parasites and their hosts. Evolution 12: 158–165. DOI 10.2307/2406026.

Mougi, A. (2012) Predator-prey coevolution driven by size selective predation can cause anti-synchronized and cryptic population dynamics. Theoretical Population Biology 81: 113–118. DOI 10.1016/j.tpb.2011.12.005.

Mougi, A. & Iwasa, Y. (2010) Evolution towards oscillation or stability in a predator-prey system. Proceedings of the Royal Society B: Biological Sciences 277: 3163–3171. DOI 10.1098/rspb.2010.0691.

Mougi, A. & Iwasa, Y. (2011) Unique coevolutionary dynamics in a predator–prey system. Journal of Theoretical Biology 277: 83–89. DOI 10.1016/j.jtbi.2011.02.015.

Northfield, T. D. & Ives, A. R. (2013) Coevolution and the effects of climate change on interacting species. Plos Biology 11: e1001685. DOI 10.1371/journal.pbio.1001685.

Nuismer, S. L. (2017) Introduction to Coevolutionary Theory. W. H. Freeman, New York, NY.

Nuismer, S. L., Doebeli, M. & Browning, D. (2005) The coevolutionary dynamics of antagonistic interactions mediated by quantitative traits with evolving variances. Evolution 59: 2073–2082. DOI 10.1111/j.0014-3820.2005.tb00918.x.

Nuismer, S. L., Ridenhour, B. J. & Oswald, B. P. (2007) Antagonistic coevolution mediated by phenotypic differences between quantitative traits. Evolution 61: 1823–1834. DOI 10.1111/j.1558-5646.2007.00158.x.

Otto, S. P. & Lenormand, T. (2002) Resolving the paradox of sex and recombination. Nature Reviews Genetics 3: 252–261. DOI 10.1038/nrg761.

Parker, M. A. (1994) Pathogens and sex in plants. Evolutionary Ecology 8: 560–584. DOI 10.1007/BF01238258.

Saloniemi, I. (1993) A coevolutionary predator-prey model with quantitative characters. The American Naturalist 141: 880–896. DOI 10.1086/285514.

Sasaki, A. & Godfray, H. C. J. (1999) A model for the coevolution of resistance and virulence in coupled host-parasitoid interactions. Proceedings of the Royal Society B: Biological Sciences 266: 455–463. DOI 10.1098/rspb.1999.0659.

Seger, J. (1988) Dynamics of some simple host-parasite models with more than two genotypes in each species. Philosophical Transactions of the Royal Society B: Biological Sciences 319: 541–555. DOI 10.1098/rstb.1988.0064.

Spottiswoode, C. N. & Stevens, M. (2012) Host-parasite arms races and rapid changes in bird egg appearance. The American Naturalist 179: 633–648. DOI 10.1086/665031.

Thompson, J. N. (2005) The Geographic Mosaic of Coevolution. The University of Chicago Press, Chicago, IL.

Tien, R. J. & Ellner, S. P. (2012) Variable cost of prey defense and coevolution in predator-prey systems. Ecological Monographs 82: 491–504. DOI 10.1890/11-2168.1.

van Velzen, E. & Gaedke, U. (2017) Disentangling eco-evolutionary dynamics of predator-prey coevolution: the case of antiphase cycles. Scientific reports 7: 17125. DOI 10.1038/s41598-017-17019-4.

Yamamichi, M. & Ellner, S. P. (2016) Antagonistic coevolution between quantitative and Mendelian traits. Proceedings of the Royal Society B: Biological Sciences 283: 20152926. DOI 10.1098/rspb.2015.2926.

Yamamichi, M. & Miner, B. E. (2015) Indirect evolutionary rescue: prey adapts, predator avoids extinction. Evolutionary Applications 8: 787–795. DOI 10.1111/eva.12295.

